# Single-stranded nucleic acid sensing and coacervation by linker histone H1

**DOI:** 10.1101/2021.03.17.435841

**Authors:** Rachel Leicher, Adewola Osunsade, Andrew P. Latham, Gabriella N. L. Chua, John W. Watters, Sophia Christodoulou-Rubalcava, Bin Zhang, Yael David, Shixin Liu

## Abstract

The linker histone H1 is the most abundant group of eukaryotic chromatin-binding proteins. The mechanism underlying the diverse physiological functions of H1 remains unclear. Here we used single-molecule fluorescence and force microscopy to observe the behavior of H1 on DNA under different tensions. Unexpectedly, we found that H1 coalesces around nascent ssDNA. Molecular dynamics simulations revealed that multivalent and transient interactions between H1 and ssDNA mediate their phase separation. We further showed that longer and unpaired nucleic acids result in more viscous, gel-like H1 droplets. Finally, we imaged H1 puncta in cells under normal and stressed conditions and observed that RPA and H1 occupy separate nuclear regions. Overall, our results provide a new perspective to understanding the role of H1 in genome organization and maintenance.

## INTRODUCTION

H1 binds the repeating unit of eukaryotic chromatin, the nucleosome core particle, by contacting the entry and exit sites of the nucleosomal DNA (*1-4*). This binding configuration underlies the well-established role of H1 in local and higher-order chromatin compaction (*5-8*). In addition, H1 functions in a variety of other DNA-templated processes including transcriptional regulation and DNA damage response (*9, 10*), the mechanisms for which are less understood. Dysregulation of H1 expression has been linked to human diseases including cancer (*11, 12*). Previous data showed that H1 is not stably associated with interphase nucleosomes (*13*) and is highly dynamic inside the nucleus (*14*). These results indicate a more complex and nuanced regulatory repertoire for H1 than a purely architectural factor. Nonetheless, dynamic H1-DNA interactions in a reconstituted biochemical setting have yet to be directly observed.

Higher eukaryotes contain multiple H1 subtypes, including eleven found in humans (*9*). Each H1 protein contains a conserved globular domain flanked by a short disordered N-terminal domain (NTD) and a long unstructured and highly charged C-terminal domain (CTD). These intrinsically disordered regions (IDRs) indicate a potential of H1 to undergo liquid-liquid phase separation (LLPS), a phenomenon that has been implicated in myriad cellular processes (*15, 16*), including chromatin organization and regulation (*17*). Indeed, recent studies reported that H1 can form condensates with double-stranded DNA (dsDNA) and nucleosomes under certain conditions (*18-21*). However, the precise mechanism and the dependence of H1-mediated phase separation on its IDRs have not been investigated.

## RESULTS

### H1 coalesces around nascent ssDNA

To directly visualize the behavior of H1 on DNA, we purified human H1.4 (hereafter referred to as H1), one of the major H1 subtypes in human cells, and labeled it with a Cy3 fluorophore (**Supplementary Fig. 1**). We then added 15 nM of Cy3-H1 to a biotinylated phage λ genomic DNA [48.5 kilobasepair (kbp)] tethered between two laser-trapped beads. By moving the beads apart from each other, we applied an increasing force to the tether and monitored H1 binding along the DNA using scanning confocal microscopy (**Fig. 1A**). At forces where the dsDNA remained in B-form, we observed a low level of H1 binding (**Fig. 1B**). As the force approached the overstretching regime (> 60 picoNewton), H1 binding markedly intensified as evidenced by a dramatic increase in total Cy3 signal across the tether (**Fig. 1C**). Strikingly, H1 specifically accumulated at both ends of the tethered DNA as shown by the emergence of two fluorescent foci (**Fig. 1B, D**; “T1”). As the inter-bead distance continued to increase, the foci became brighter, migrating towards each other, and eventually merging into one singular spot (**Fig. 1B, D**; “T2”). Because the DNA was attached to the bead via the 3’ end of each strand, we posited that H1 preferentially binds to the untethered single-stranded DNA (ssDNA) created by force-induced unpeeling (**Fig. 1E**). Notably, H1 did not bind the other ssDNA strand that was attached to the bead (i.e. we detected no fluorescence signal between the H1 focus and its proximal bead) (**Fig. 1B, E**). Hence, under these experimental conditions, H1 does not bind ssDNA under tension. In other examples, we also observed H1 foci forming internally, presumably around relaxed ssDNA originating from the internal nicks naturally occurring within the DNA substrate (**Supplementary Fig. 2**). In addition, we found this behavior to be unique to the linker histone, since core histones such as H2B exhibited force-insensitive binding and did not coalesce on ssDNA created by high tension (**Supplementary Fig. 3**). This difference may be attributed to the fact that H1 proteins are structurally more disordered and have a higher net positive charge than the core histones (*9*).

**Figure 1.**
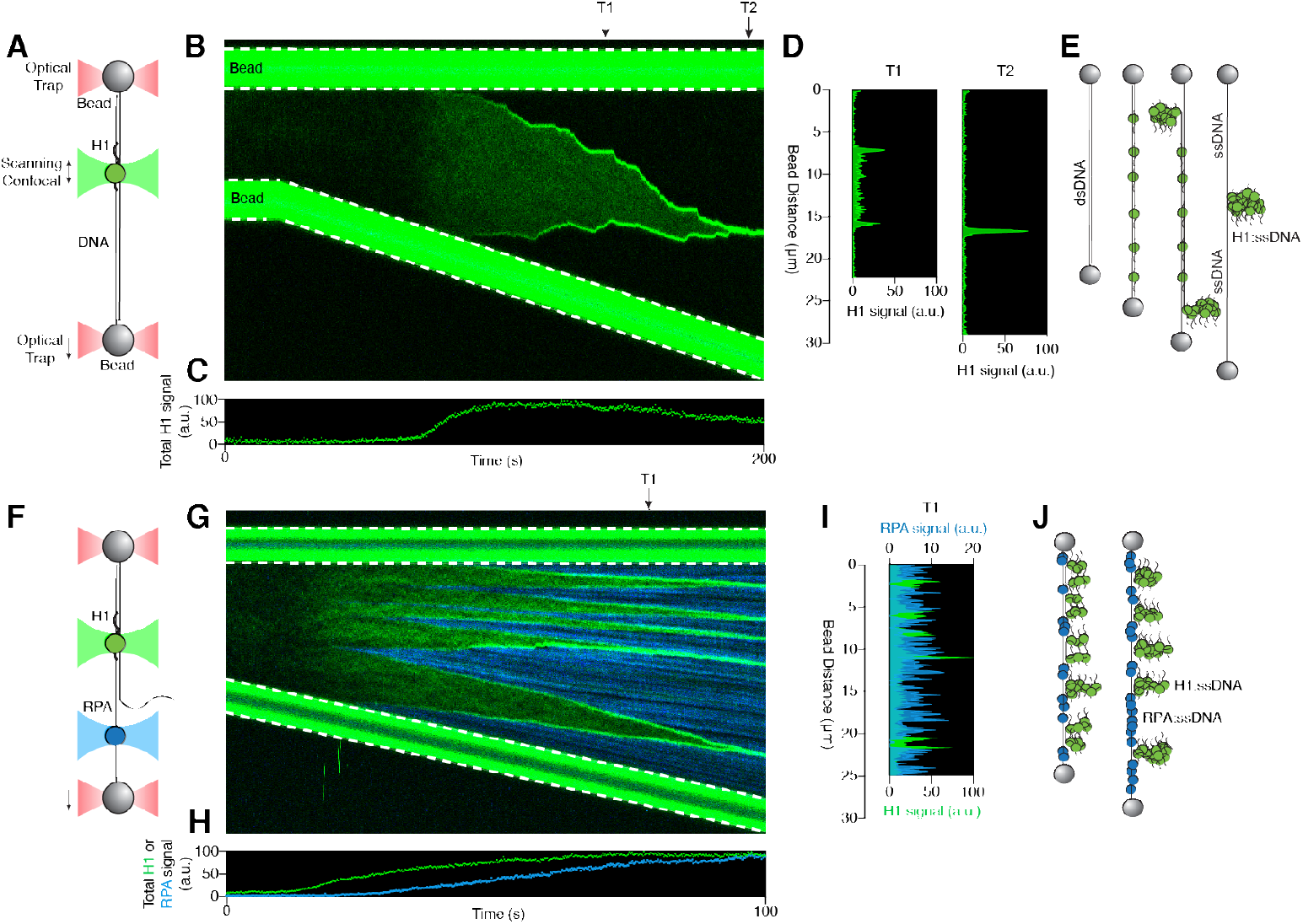
H1 coalesces around nascent ssDNA. (**A**) Schematic of the combined single-molecule fluorescence and force microscopy. A biotinylated λ-DNA molecule (48.5 kbp) is tethered between two streptavidin-coated polystyrene beads. (**B**) A representative kymograph of Cy3-H1 binding to DNA over time as the inter-bead distance was increased. (**C**) Total H1 signal across the DNA as a function of time for the kymograph shown in (B). (**D**) Distribution of the H1 signal along the DNA at two specific time points (T1 and T2) as indicated by the arrows in (B). (**E**) Cartoon illustrating the distinct binding configurations of H1 on DNA under different tensions. ssDNA is created by force-induced unpeeling. (**F**) Schematic of two-color imaging for simultaneous visualization of H1 and RPA binding to DNA. (**G**) A representative kymograph of Cy3-H1 (green) and AlexaFluor488-RPA (blue) binding to DNA over time as the inter-bead distance was increased. (**H**) Total H1 and RPA signals across the DNA as a function of time for the kymograph shown in (G). (**I**) Distribution of the H1 (green) and RPA (blue) signals along the DNA at a specific time point (T1) as indicated by the arrow in (G). (**J**) Cartoon illustrating that H1 and RPA occupy separate regions of the tethered DNA. H1 coalesces around relaxed ssDNA, whereas RPA binds to ssDNA under tension.

**Figure 2.**
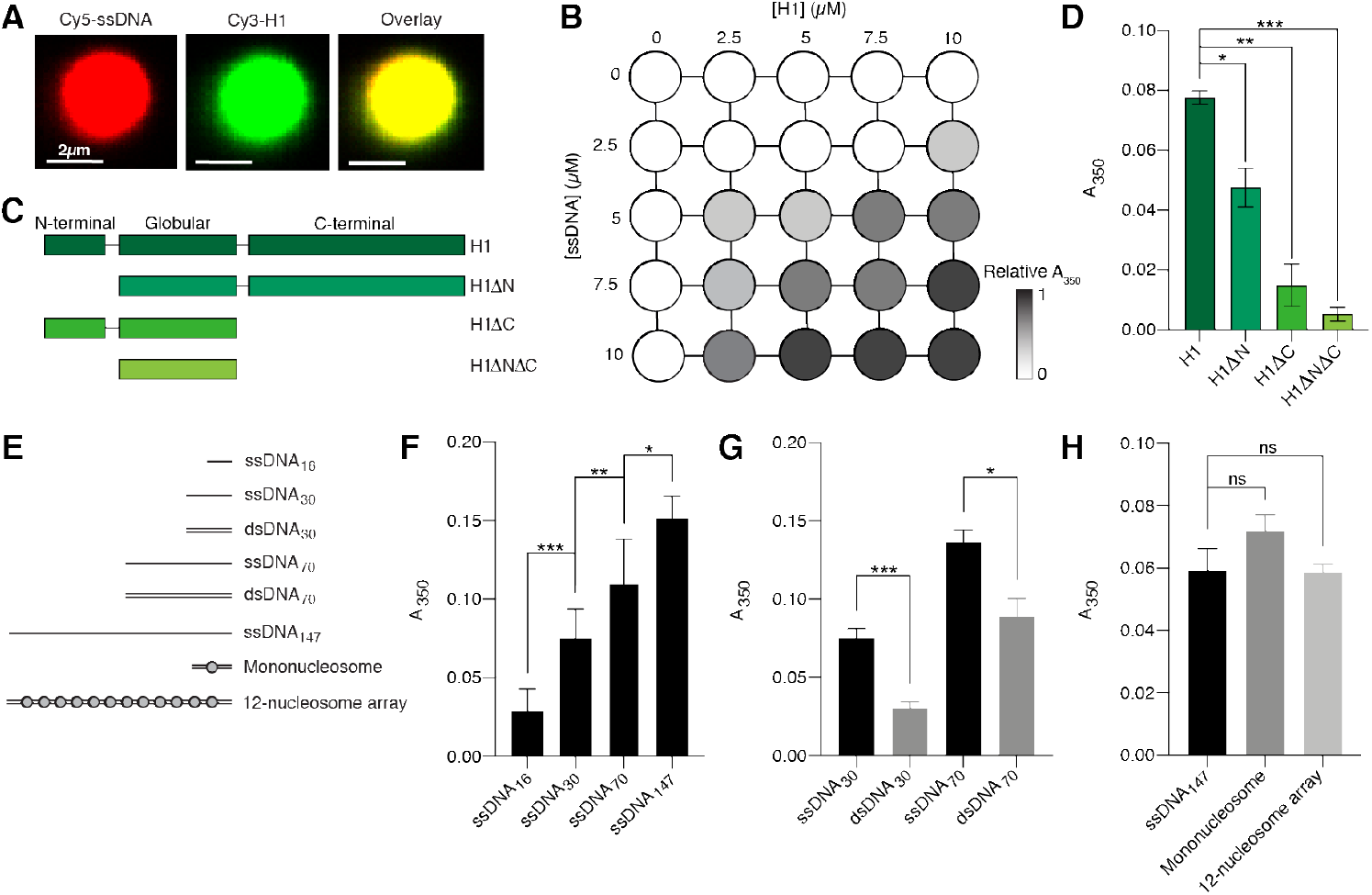
H1 phase separation depends on its disordered tails and discriminates between ssDNA versus dsDNA. (**A**) Fluorescence images of a representative droplet formed by mixing 2.5 µM Cy3-H1 with 10 µM Cy5-ssDNA_75_ (10% labeled). (**B**) Matrix diagram of solution turbidity values (A_350_) measured at different concentrations of H1 and ssDNA_75_ (both unlabeled), normalized by the highest A_350_ value at 10 µM H1 and 10 µM ssDNA. (**C**) Schematic of the domain structures of full-length H1 and NTD/CTD-truncated H1 constructs. (**D**) A_350_ values for different H1 constructs depicted in (C) mixed with 10 µM ssDNA_75_. (**E**) Schematic of the DNA and chromatin substrates used in the phase separation experiments with H1. (**F**) A_350_ values for full-length H1 mixed with ssDNA of different lengths (44 µM ssDNA_16_, 23 µM ssDNA_30_, 10 µM ssDNA_70_, and 4.8 µM ssDNA_147_). ssDNA concentrations were normalized to yield the same total amount of nucleotides. (**G**) A_350_ values for H1 mixed with ssDNA or dsDNA of the same length and sequence (ssDNA_30_/dsDNA_30_, ssDNA_70_/dsDNA_70_, all in 10 µM). (**H**) A_350_ values for H1 mixed with ssDNA_147_ (5 µM), mononucleosomes (5 µM), or 12-nucleosome arrays (600 nM). All experiments were performed with 2.5 µM H1 unless specified otherwise. Data are mean ± SEM of at least three independent measurements.

**Figure 3.**
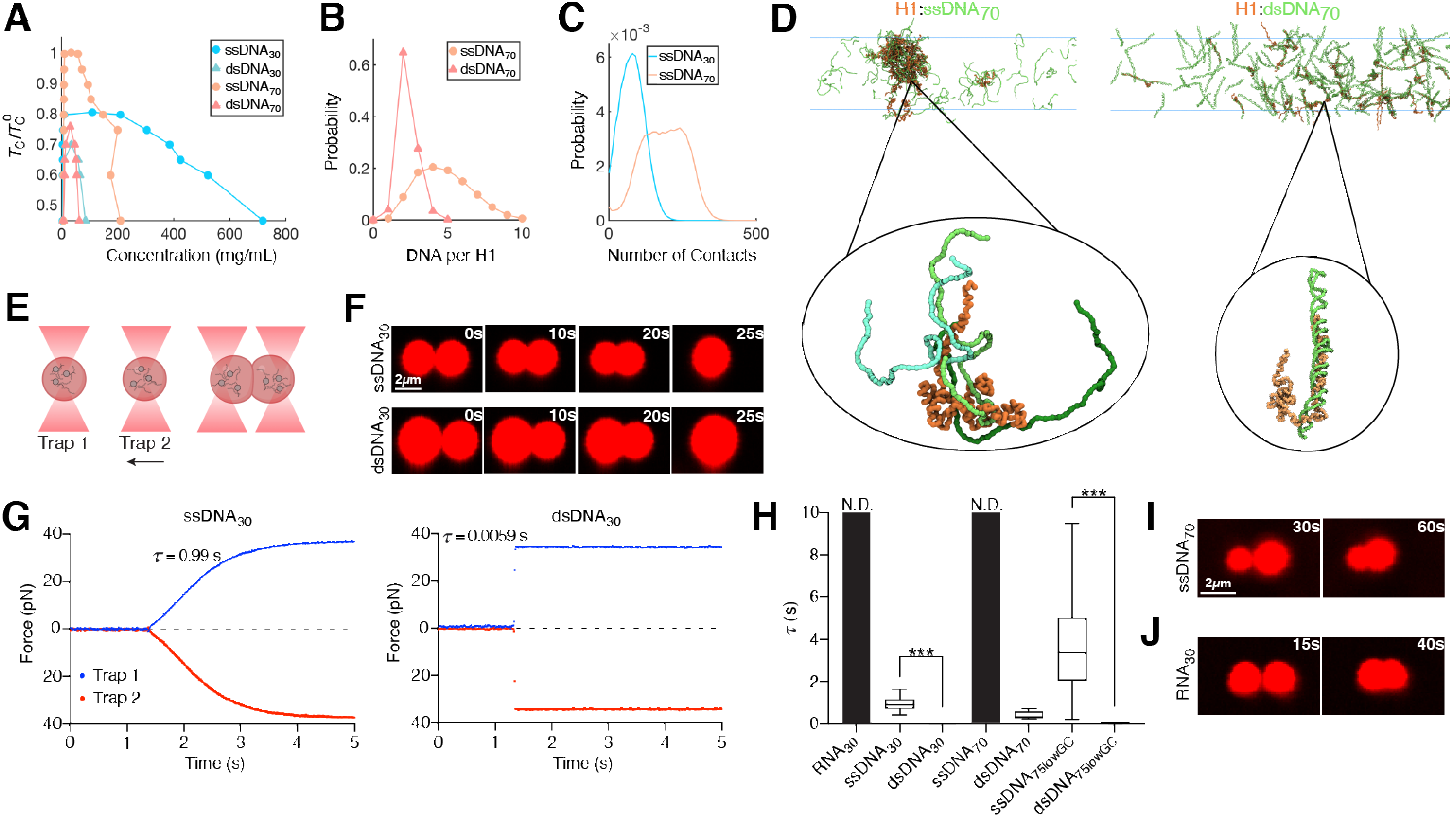
H1:ssDNA and H1:dsDNA droplets exhibit distinct material properties. (**A**) Computational phase diagrams in the temperature-concentration plane. For each system, we calculated the upper critical temperature for phase separation (*T*_C_). The displayed values are normalized by the *T*_C_ with ssDNA_70_ (*T*_C_^0^). (**B**) Probability distribution of the number of DNA bound to each H1. (**C**) Probability distribution of the number of contacts with H1 formed by an individual DNA molecule (ssDNA_30_ or ssDNA_70_). (**D**) Example configurations of the condensates formed between ssDNA_70_ (left) and dsDNA_70_ (right) with H1. The insets highlight the different protein-DNA binding modes. (**E**) Schematic of the controlled droplet fusion assay with optical tweezers (droplet created with BioRender). (**F**) Snapshots of a representative fusion event for H1:Cy5-ssDNA_30_ (top) and H1:Cy5-dsDNA_30_ (bottom) droplets visualized by Cy5 fluorescence. (**G**) Force profiles of the two traps during a representative fusion event for H1:ssDNA_30_ (left) and H1:dsDNA_30_ (right) droplets. *τ* represents the droplet fusion time. (**H**) *τ* values for H1 droplets formed with different nucleic acid substrates. The *τ* value for dsDNA_30_ is too small to be visible (0.0046 ± 0.0013 s). *τ* values for RNA_30_ and ssDNA_70_ were nondetermined (N.D.) because these droplets did not fuse during our observation window (at least 20 s after contact). Data are mean ± SEM of at least 12 fusion events for each condition. (**I** and **J**) Representative images of two H1:Cy5-ssDNA_70_ droplets or two H1:Cy5-RNA_30_ droplets that were unable to fuse. All experiments were performed with 2.5 µM H1 and 10 µM DNA/RNA (10% labeled).

To confirm this interpretation, we used AlexaFluor488-labeled Replication Protein A (RPA), a well-studied eukaryotic ssDNA-binding protein that is capable of interacting with ssDNA under tension (**Fig. 1F**). Indeed, both H1 and RPA signals on DNA increased with tension (**Fig. 1G, H**), but with drastically distinct localization patterns. H1 foci were consistently observed on the flanks of RPA-bound regions of the tether in an anticorrelated manner (**Fig. 1G, I**). This finding suggests that H1 does not bind to ssDNA regions that are under tension, whereas RPA is excluded from H1 foci around the relaxed ssDNA (**Fig. 1J**). Next we examined whether the formation of H1 foci is reversible by sequentially stretching and relaxing the tether. Interestingly, we found that the majority of H1 foci dissolved upon tether relaxation (**Supplementary Fig. 4A**), presumably induced by the energetically favorable re-annealing of the two complementary ssDNA. In contrast, in the presence of RPA, most of the H1 foci persisted at low forces (**Supplementary Fig. 4B, C**), suggesting that RPA poses a barrier against the re-annealing of ssDNA and the dissolution of H1 foci. Overall, these results reveal that H1 preferentially binds to and coalesces around relaxed ssDNA, forming condensate-like complexes.

**Figure 4.**
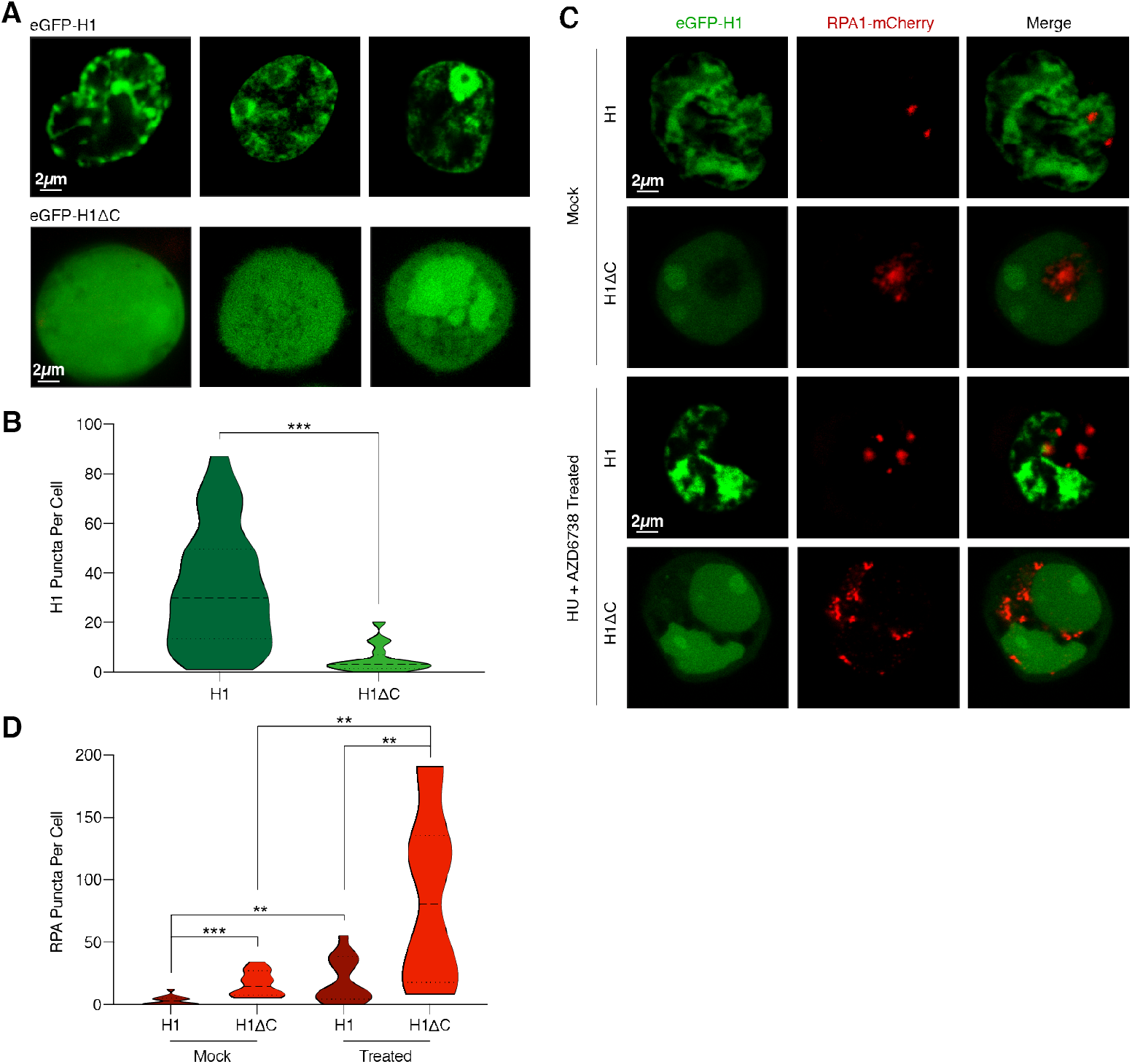
H1 puncta formation in the cell requires its C-terminal tail. (**A**) Representative confocal fluorescence images of HEK293T cells transfected with either eGFP-H1 (top) or eGFP-H1Δ C (bottom). (**B**) Violin plot showing the distribution of number of H1 puncta per cell for eGFP-H1 and eGFP-H1Δ C cells. (**C**) Representative images of cells co-transfected with eGFP-H1 and RPA-mCherry, or with eGFP-H1Δ C and RPA-mCherry, and treated with either mock or 2 mM HU + 20 µM AZD6738 for 12 h. (**D**) Violin plot showing the distribution of number of RPA puncta per cell for eGFP-H1 and eGFP-H1Δ C cells after mock or HU + AZD6738 treatment.

### Phase separation of H1 with ssDNA depends on its disordered tails

Based on the single-molecule observations, we hypothesized that H1 forms phase-separated condensates with ssDNA. To test this hypothesis, we mixed full-length H1 with ssDNA of 75 nucleotide (nt) in length under physiological buffer conditions, which indeed yielded distinct phase-separated droplets (**Supplementary Fig. 5A**). We further validated the enrichment of H1 and ssDNA within the same droplets via fluorescence imaging using Cy3-labeled H1 and Cy5-labeled ssDNA (**Fig. 2A**). The observed LLPS propensity increased as a function of H1 and ssDNA concentrations, as shown by solution turbidity measurements (absorbance at 350 nm, A_350_) (**Fig. 2B**). Droplet formation required the presence of DNA, but not the other buffer components such as polyethylene glycol (PEG), nor was it affected by the cognate H1 chaperone prothymosin *α* (ProT*α*) that was shown to form a high-affinity complex with H1 *in vitro* while remaining disordered (*24*) (**Supplementary Fig. 5A-C**).

To determine the effect of the IDRs of H1 on its LLPS behavior, we truncated either the NTD or CTD or both (H1Δ N, H1Δ C, H1Δ NΔ C, respectively) and assessed the capacity of these truncated H1 proteins to phase separate with ssDNA (**Fig. 2C**). We found that H1:ssDNA LLPS is significantly impaired by deletion of the longer CTD and to a lesser extent, deletion of the NTD (**Fig. 2D**). When both tails were removed, the globular domain alone showed minimal phase separation with ssDNA (**Fig. 2D**). These results experimentally prove the expectation that the LLPS ability of H1 is mediated by its unstructured tails.

### Tight binding of H1 with dsDNA suppresses phase separation

Next, we examined how different properties of the DNA may impact its tendency to phase separate with H1. Using ssDNA of different lengths ranging from 16 to 147 nt (**Fig. 2E**), we found that longer ssDNA promotes H1:ssDNA droplet formation after, importantly, normalizing for the total amount of nucleotides (**Fig. 2F**). Then we compared pairs of ssDNA and dsDNA of 30 or 70 nt/bp in length (ssDNA_30_ vs. dsDNA_30_; ssDNA_70_ vs. dsDNA_70_). A_350_ measurements showed that ssDNA possesses a significantly higher capacity to phase separate with H1 compared with dsDNA of the same length and sequence (**Fig. 2G**), in accordance with the single-molecule results shown above. This difference is even more pronounced when we normalized the DNA concentrations to the total number of nucleotides between ssDNA and dsDNA (**Supplementary Fig. 5D**). In addition, we examined chromatinized DNA (**Fig. 2E**), which was shown to phase separate with H1 (*19, 20*), and found that bare ssDNA possesses a similar ability to form droplets with H1 compared to mononucleosomes or nucleosome arrays (**Fig. 2H**).

To obtain a detailed structural characterization of H1:DNA condensates, we carried out coarse-grained molecular dynamics (MD) simulations (*25, 26*). The force fields used in these simulations were fine-tuned to capture sequence-specific protein-protein interactions and the persistence lengths of ssDNA and dsDNA. The simulated critical temperatures (*T*_c_) of phase separation in different systems match the A_350_ trends recorded experimentally (**Fig. 3A, Supplementary Fig. 6A, B**), supporting the usefulness of the *in silico* model for mechanistic exploration. Close examination of the simulated condensed phases reveals marked differences in their internal organization. H1 molecules coacervate with ssDNA into droplets at much higher densities than with dsDNA (**Fig. 3B-D, Supplementary Fig. 6D**). On the other hand, dsDNA molecules are more rigid, rendering them more resistant to dense packing (**Fig. 3D**).

Simulation results suggest that H1 adopts distinct binding modes depending on the type and length of DNA. H1 coils around dsDNA to achieve tight binding, while the interactions between H1 and ssDNA are more transient (**Fig. 3D**). The fewer contacts with individual ssDNA allow each H1 to simultaneously contact multiple ssDNA molecules (**Fig. 3B**). Such multivalent interactions could explain ssDNA’s increased propensity for phase separation (*27*). Furthermore, longer DNA binds H1 more tightly by forming more contacts with H1 (**Fig. 3C**), yielding lower overall diffusion coefficients (*D*) of H1 (**Supplementary Fig. 6C**). The relative affinity between binding partners is another known factor that dictates droplet properties (*28*). Thus, the strength and multivalency of H1:DNA interactions collectively determine the observed phase separation trends. Lastly, in agreement with our experimental observations that the disordered tails of H1 control its phase separation, simulations also show that H1 interacts with DNA primarily through its CTD, although the NTD also contributes significantly (**Supplementary Fig. 6E, F**).

### H1:ssDNA and H1:dsDNA droplets exhibit distinct material properties

To experimentally dissect the difference in physicochemical properties between H1:ssDNA and H1:dsDNA droplets, we leveraged the ability of optical tweezers to trap and manipulate protein droplets due to their different refractive index relative to the surrounding medium (*29-31*). We first generated H1:ssDNA droplets containing fluorescently labeled ssDNA for real-time visualization. Two micron-sized droplets were captured and brought into proximity by moving one trap towards the other (**Fig. 3E**). We observed that the two droplets eventually fused into a singular one after contact (**Fig. 3F**). Meanwhile, the force experienced by the droplets was recorded during the fusion process, displaying a gradual increase as the droplets were being pulled away from their respective trap center (**Fig. 3G, Supplementary Fig. 7**). When fusion was completed, the merged droplet was caught between two traps, yielding a steady-state force readout. This force measurement allowed us to quantify the timescale of the fusion process (*τ*), which lasted 0.96 ± 0.37 s for the H1:ssDNA_30_ droplets (**Fig. 3H**). We next performed the same droplet fusion experiment with H1:dsDNA_30_ droplets (**Fig. 3F**). We noticed that dsDNA droplets fused substantially faster (0.0046 ± 0.0013 s) than ssDNA ones (**Fig. 3G, H, Supplementary Fig. 7**). The ∼200-fold difference in the fusion kinetics suggests that H1:ssDNA droplets are more viscous and gel-like, whereas H1:dsDNA droplets are more liquid-like, which is corroborated by our MD simulation results. In addition, we conducted fluorescence recovery after photobleaching (FRAP) experiments, which provides information about the fluidity of the droplets. We found that the recovery rate for H1:dsDNA droplets was faster than that for H1:ssDNA ones, again suggesting a more dynamic and liquid-like droplet (**Supplementary Fig. 8**).

Next we used the droplet fusion assay to evaluate how DNA length and base composition affect the material properties of H1 condensates. We found that a longer ssDNA_70_ with the same GC content as ssDNA_30_ renders the droplets unable to fuse over the observation window (**Fig. 3I**). Conversely, lowering the GC content of the ssDNA fluidifies the droplets and accelerates the fusion process [compare ssDNA_70_ (50% GC) and ssDNA_75_ (33% GC) in **Fig. 3H**]. Deletion of the NTD of H1 also makes the droplets more liquid-like, as evidenced by a shortened fusion time and faster FRAP recovery for H1Δ N-containing droplets (**Supplementary Fig. 8B, C**). Deletion of the H1 CTD completely abolished droplet formation, therefore no measurements could be made. Notably, we observed that H1 also formed condensates with single-stranded RNA. However, compared to ssDNA of the same length and sequence, H1:RNA droplets took longer to fuse, if at all (**Fig. 3H, J**), suggesting that H1 differentially coacervates with DNA versus RNA.

### H1 C-terminal domain mediates puncta formation in cells

Finally, to observe the H1 localization pattern in a native context, we expressed H1.4 tagged with eGFP (eGFP-H1) in HEK293T cells and performed live-cell confocal imaging. Cells expressing full-length eGFP-H1 displayed multiple nuclear puncta (**Fig. 4A**), consistent with previously observed patterns (*20*). In contrast, cells expressing eGFP-tagged CTD-deleted H1 (eGFP-H1Δ C) displayed a striking deficiency in puncta formation, instead exhibiting a diffused signal throughout the nucleus (**Fig. 4A, B**). Of note, we verified that recombinant eGFP-tagged H1 can still phase separate with ssDNA, and that eGFP-H1:ssDNA droplets can fuse *in vitro* (**Supplementary Fig. 9**).

To better understand the relationship between ssDNA and H1 nuclear puncta, we co-transfected cells with eGFP-H1 and RPA1 tagged with mCherry (RPA-mCherry). In agreement with our *in vitro* single-molecule results, H1 and RPA localization appeared to be mutually exclusive (**Fig. 4C**). Notably, eGFP-H1Δ C cells contained significantly more RPA puncta than eGFP-H1 cells (**Fig. 4C, D**), suggesting that the depletion of H1:ssDNA condensates frees up additional ssDNA sites for RPA to bind. We next treated cells with both hydroxyurea (HU) and the ATR kinase inhibitor ceralasertib (AZD6738). HU induces replication stress and the accumulation of ssDNA in the cell, while AZD6738 impairs the DNA damage response (*32, 33*). Following treatment, we detected a significant increase in the number of RPA puncta in both eGFP-H1 and eGFP-H1Δ C cells (**Fig. 4C, D**). Interestingly, the number of H1 puncta also increased in eGFP-H1 cells (**Supplementary Fig. 10**). These results show that the CTD of H1 is critical for H1 puncta formation *in vivo* and that these puncta exclude RPA.

## DISCUSSION

Nucleic acids, mainly RNA, are known to modulate the viscoelastic properties of protein droplets (*34, 35*). Recently, differential phase separation with ss versus ds DNA has been observed for model peptides and proteins (*36-38*). Here we report that H1, a prominent and highly abundant protein in eukaryotic cells, preferentially interacts with ssDNA and forms condensates with material properties modulated by the ssDNA length and sequence. Our *in silico* modeling suggests that the tighter binding of H1 to dsDNA than to ssDNA results in a lower phase separation propensity. While affinity above a certain threshold is required for LLPS, very stable contacts—in which two polymers can only interact in a one-to-one fashion—turn out to suppress the multivalent interactions needed for phase separation (*39*). The differential condensation of H1 with ssDNA versus dsDNA is likely dependent on the relative abundance of these species in the nucleus, which fluctuates during the cell cycle. In circumstances where ssDNA accumulates, such as during DNA replication and repair, H1:ssDNA condensation may become a significant mechanism for facile and reversible regulation of DNA metabolism.

The ability of H1 to sense different forms of nucleic acids, and potentially partition them into distinct compartments, implies an expanded role for H1 in local and global genome organization and maintenance, beyond its known function in chromatin compaction. Our results *in vitro* and in cells suggest that H1 and RPA compete for binding to ssDNA. H1-induced ssDNA coacervation may allow for ssDNA sequestration and exclusion of certain DNA-processing enzymes. In accordance with this view, H1 has been shown to inhibit homologous recombination, and its depletion leads to altered sensitivity to DNA damage (*40, 41*). Based on our single-molecule data, while H1 occupies the majority of ssDNA under slack, only RPA can interact with ssDNA under tension, which could be exerted by force-generating motor proteins in the cell, thereby activating RPA-mediated reaction pathways. It is noteworthy that H1:RNA droplets are distinguished by their gel- or solid-like behavior. Given the prevalence of both coding and non-coding RNAs in the cell, H1:RNA condensates may have important biological implications.

Human H1 subtypes diverge in homology primarily in their CTD composition. While these variants exhibit distinct abilities to bind and compact chromatin (*42*), the extent of this functional variability remains controversial. Our results provide direct evidence that the CTD is a key mediator of H1 condensation with nucleic acids. It is therefore possible that the divergent CTDs of H1 subtypes may enhance or attenuate their ability to phase separate, resulting in distinct roles for each subtype in genome organization. The experimental and computational platforms developed in this study pave the way for a systematic interrogation of H1 subtypes, posttranslational modifications, and disease-associated mutations, which together constitute a highly regulated and multifaceted network of linker histones (*43*).

These efforts may explain how, for example, H1 can instigate both global changes in chromatin structure as well as specific alterations in gene expression (*44*).

## METHODS

### Oligo sequences

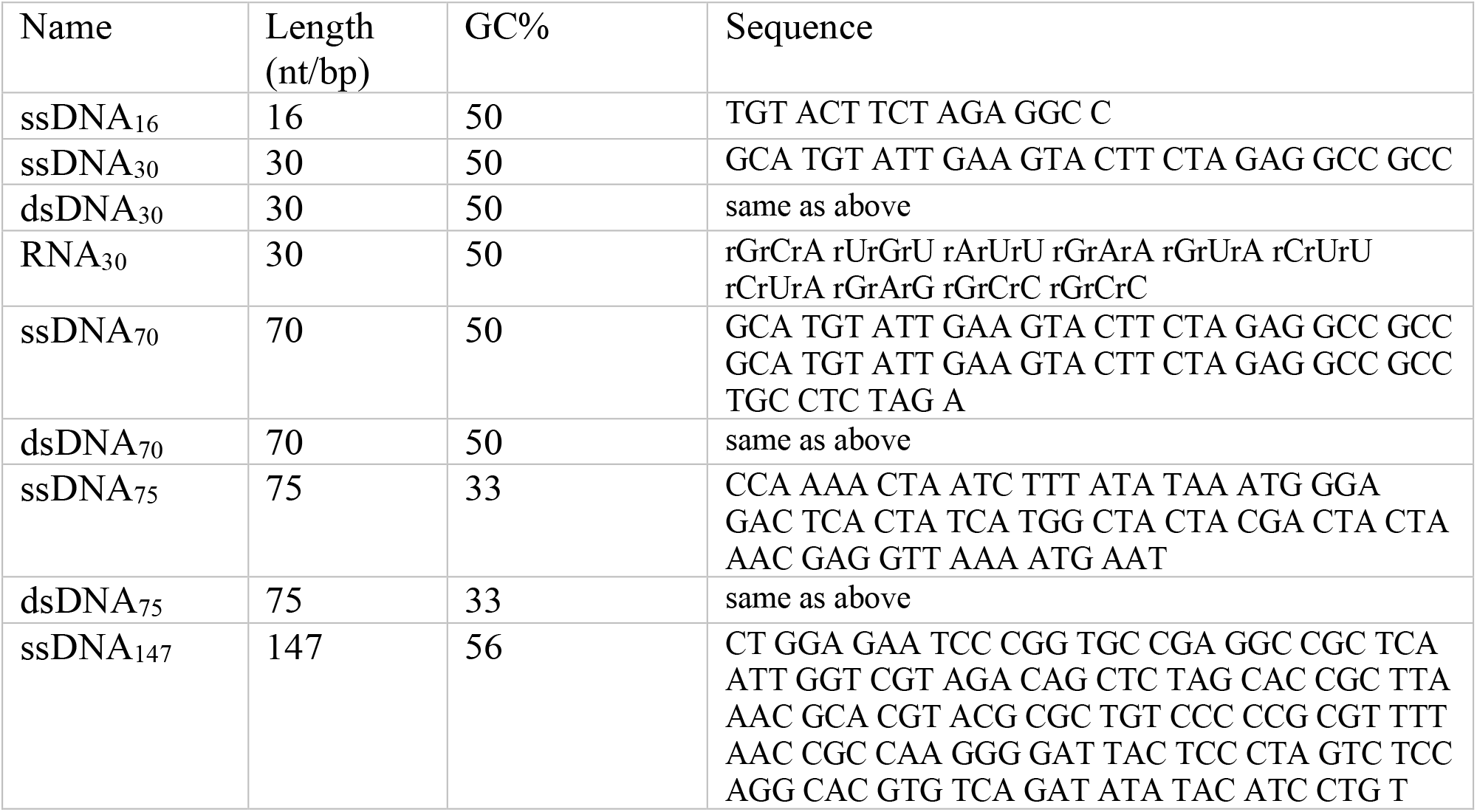

### Protein purification and labeling

All recombinant H1 proteins and mutants were purified as described previously with minor modifications (*42*). Briefly, Rosetta DE3 cells expressing His-SUMO-H1.4-GyrA-His were induced with IPTG and expressed for 12 hours before harvesting. After rod sonication, lysate was incubated with Ni-NTA beads (Bio-Rad) and eluted. The eluent was treated with recombinant Ulp-1 (1:100 w/v) and 500 mM β-mercaptoethanol, followed by the addition of solid urea to a final concentration of 6 M and the pH was adjusted to 9.0. This was then loaded onto a HiTrap SP cation exchange column (Cytiva) and subjected to a gradient of 100% H1 purification buffer A (6 M urea, 20 mM Tris pH 9.0, 200 mM NaCl) to 100% buffer B (6 M urea, 20 mM Tris pH 9.0, 1 M NaCl) of an AKTA FPLC system (GE Healthcare). Fractions containing full-length H1 were pooled and purified on a semi-preparative C18 HPLC column on a gradient of 0—70% buffer B and freeze-dried until use. H1 proteins were resuspended in H1 buffer (20 mM Tris pH 7.5, 200 mM NaCl) before use.

Recombinant H1.4A4C was purified as described above. Fluorescent labeling was performed based on a previously published protocol (*45*). Briefly, H1.4A4C was dissolved in 4 mL of histone labeling buffer (6 M Guanidine, 20 mM Tris pH 7.5, 0.2 mM TCEP). Three molar equivalents of Cy3-maleimide (ApexBio) in DMF were added and mixed gently at room temperature followed by the addition of 1 mM β-mercaptoethanol to quench the reaction. The resultant mixture was purified on a semi-preparative reverse-phase C18 HPLC column on a gradient of 0—70% buffer B, and freeze-dried before resuspension in H1 buffer. Recombinant human H2B T49C was purified and labeled as previously described (*46*). AlexaFluor488-RPA was prepared as described previously (*47*).

### Correlative single-molecule fluorescence and force assay of H1-DNA interactions

#### DNA preparation

To create a terminally biotinylated dsDNA template, the 12-base 5’ overhang on each end of genomic DNA from bacteriophage λ (48,502 bp; Roche) was filled in with a mixture of natural and biotinylated nucleotides by the exonuclease-deficient DNA polymerase I Klenow fragment (New England BioLabs). Reaction was conducted by incubating 10 nM λ-DNA, 33 μM each of dGTP/dATP/biotin-11-dUTP/biotin-14-dCTP (Thermo Fisher), and 5 U Klenow in 1× NEB2 buffer at 37 °C for 45 min, followed by heat inactivation for 20 min at 75 °C. DNA was then ethanol precipitated overnight at −20 °C in 2.5× volume cold ethanol and 300 mM sodium acetate pH 5.2. Precipitated DNA was recovered by centrifugation at 20,000× g for 15 min at 4 °C. After removing the supernatant, the pellet was air-dried, resuspended in TE buffer (10 mM Tris-HCl pH 8.0, 1 mM EDTA) and stored at 4 °C.

### Single-molecule experiments

Single-molecule experiments were performed at room temperature on a LUMICKS C-Trap instrument (*47*). A computer-controlled stage enabled rapid movement of the optical traps within a microfluidic flow cell. Laminar flow separated channels 1—3, which were used to form DNA tethers between 3.23-μm streptavidin-coated polystyrene beads (Spherotech) held in traps with a stiffness of 0.6 pN/nm. Under constant flow, a single bead was caught in each trap in channel 1. The traps were then quickly moved to channel 2 containing the biotinylated DNA. By moving one trap against the direction of flow but toward the other trap, and vice versa, a DNA tether could be formed and detected via a change in the force-extension (*F*-*x*) curve. The traps were then moved to channel 3 containing only buffer, and the presence of a single DNA was verified by the *F*-*x* curve. Orthogonal channel 4 served as a protein loading and imaging channel. Flow was turned off during data acquisition. Force data were collected at 100 kHz. AlexaFluor488 and Cy3 fluorophores were excited by 488-nm and 532-nm laser lines, respectively. Kymographs were generated via a confocal line scan through the center of the two beads.

For H1 and RPA binding experiments, single DNA tethers were moved to channel 4 where they were incubated with 15 nM Cy3-H1. DNA was stretched between the beads by moving the right trap at a constant velocity (0.1 µm/s) in the *x*-direction. In experiments with H1 and RPA, 10 nM AlexaFluor488-RPA was added simultaneously to channel 4. In experiments with H2B, single DNA tethers were incubated with 15 nM Cy3-H2B and imaged as described for H1.

### Data analysis

Bead photon counts were removed from the kymographs using either manual removal or a local search method to maximize the sum over the size of the beads’ autofluorescence. The distance and confocal photon count measurements between these removed regions were then used to plot line scans of the kymographs. A photon count threshold value was used to define the foci points in the kymograph traces (110% of the maximum photon count from the dsDNA region of the kymograph). To analyze the reversibility of H1 foci formation, the lumicks.pylake Python package’s greedy line tracking algorithm was applied to define line traces in the regions where the DNA tether was being relaxed (*48, 49*). Line traces present in the first 20% of the region were analyzed. A trace was counted as dissolved if the trace ended before the last 20% of the region. Traces that were still present in the last 20% of the region were counted as retained.

### Solution turbidity measurements

Phase separation experiments were performed in 20-µL volumes of 20 mM Tris-HCl pH 7.5, 200 mM NaCl, and 10% PEG8000, at a DNA concentration of 10 µM and an H1 concentration of 2.5 µM unless otherwise specified. Absorbance measurements were taken at 350 nm on a NanoDrop spectrophotometer (Thermo Scientific) after 10-min incubation.

### Droplet imaging and manipulation

#### Droplet imaging

H1:nucleic acid droplets were formed as described above for the solution turbidity assay and injected into a home-made flow cell. Droplets were imaged on the C-Trap instrument either by brightfield microscopy or fluorescence microscopy using a 532-nm or 639-nm laser. For fluorescence recovery after photobleaching (FRAP) experiments, droplets were partially bleached (100% laser power) for 2 s, followed by 2D confocal scanning (10% laser power) every 5 s using a custom script titled “FRAP droplet imaging” (https://harbor.lumicks.com/single-script/3a796fac-dbb3-4fe1-8ce7-8b0cf8c25ad9). Images were uploaded to FIJI as a stack and analyzed by the FRAP Profiler plugin.

#### Controlled droplet fusion experiments

Two droplets were captured by the 1064-nm infrared laser in the dual traps of C-Trap (5% laser power) and visualized by 2D confocal scanning every 5 s. One trap was manually stepped towards the other trap in 200-nm intervals after every image scan until the fusion process started. The force for each trap was recorded concurrently.

#### Data analysis

The droplet fusion times (*τ*) were calculated by fitting sigmoidal curves to the force-time data over the time windows of droplet fusion. The fit equation used is 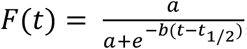, where *F* is the normalized magnitude of high-frequency force data (78 kHz) from the stationary optical trap, *t* is the time value of each force data point, *a* and *b* are the generalized fit parameters, and *t*_1/2_ is another fit parameter that approximates the time of half-maximum force. The fits were then used to calculate *τ* as defined by the time lag between 30% and 80% of the normalized force values.

Force and fluorescence data from the .h5 files generated from C-trap experiments were analyzed using tools in the lumicks.pylake Python library supplemented with other Python modules (Numpy, Matplotlib, Pandas) (*50*) in a custom GUI Python script titled “C-Trap .h5 File Visualization GUI” (https://harbor.lumicks.com/single-script/c5b103a4-0804-4b06-95d3-20a08d65768f). This script was used to extract confocal images and fusion traces from droplet formation, fusion, and FRAP experiments.

### MD Simulations

#### Force field

Coarse-grained, implicit-solvent molecular dynamics simulations were performed with one bead per protein residue or DNA base using the maximum entropy optimized force field (MOFF) for proteins (*26*) and the molecular renormalization group coarse-graining model (MRG-CG) for DNA (*25*). The MOFF force field was complemented with structure-based modeling potentials in the ordered domain of H1 to stabilize its tertiary structure (*51*). The strength of the structure-based potentials was tuned to reproduce the root-mean-square fluctuations from all-atom simulations. The original DNA model was parameterized with explicit ions. We rescaled its bonded, angle, and fan interactions by a factor of 0.9 to reproduce the persistence lengths of ssDNA and dsDNA in simulations with implicit ions. Electrostatic interactions within and between protein and DNA molecules were described using the Debye-Huckel potential and a distance-dependent dielectric constant (*26*). A salt concentration of 150 mM was used in simulations to account for the uneven partition of salts in complex coacervation, with lower values in the condensed phase (*52*). In addition to electrostatic interactions, an excluded volume term of 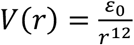 was introduced to prevent overlap between protein and DNA molecules, where *r* is the distance between two beads and *ϵ*_0_ = 1.6264 × 10^−3^ kJ mol^-1^ nm^12^. All simulations were performed using the GROMACS simulation package (*53*).

#### Slab simulation methodology

Simulations with a ratio of 4DNA:1H1 were performed following the slab methodology to determine critical temperatures (*26, 54*). The ssDNA simulations included 160 DNA molecules and 40 H1 molecules, while those with dsDNA were performed with 320 DNA molecules and 80 H1 molecules. To check for finite-size effects, we also performed simulations with 160 DNA molecules and 40 H1 molecules for dsDNA and obtained similar results. For each setup, the molecules were initially placed in a large simulation box of size 100 nm × 100 nm × 100 nm. We then performed the steepest descent energy minimization, followed by a 0.1 µs long NPT simulation at 150 K and 1 bar using a Parrinello-Rahman isotropic barostat and time coupling constant of 1 ps. The NPT simulation collapsed the molecules into a dense phase. The *z*-dimension of the simulation box was then expanded by ∼20 times the original size, resulting in a droplet with a dilute phase on either side. We then performed an NVT simulation for 0.1 µs with a time coupling constant of 100 ps. During the simulation, we raised the temperature from 150 K to the desired value. The resulting equilibrated system was run for 2 µs in the NVT ensemble. Configurations collected at every 1-ns steps from the second half of the trajectories were used for analysis.

#### Data analysis

For each simulated configuration, we first computed a molecular contact matrix. Contacts between two molecules, protein or DNA, were determined if any of their particles are within 1 nm. Using a depth-first search algorithm over the network constructed from the contact matrix, we identified the largest cluster. The high and low densities at a given temperature were determined using molecules whose centers of mass were in regions within 2.5 nm of the largest cluster or 150 nm or more away from it. These densities were used to obtain the critical temperature (*T*_C_) by fitting to the analytical expression *ρ*_*H*_ −*ρ*_*L*_ = *A*(*T*_*C*_ − *T*)^*β*^, where *β* = 0.325 is the universality class of a 3D Ising model. Unlike the analytical Ising model, which assumes a homogeneous system, our system is highly heterogeneous, with strong H1:DNA interactions and substantially weaker H1:H1 and DNA:DNA interactions. As such, we note that there is additional uncertainty in *T*_C_ arising from the sensitivity in density estimation. In particular, while *T*_C_ was predicted to be less than 300 K for some systems, large clusters with a significant percentage of H1 are prevalent. Therefore, we used simulation data obtained at 300 K to analyze protein-DNA binding and diffusion coefficients. MDAnalysis was used to help with analysis (*55*).

#### Live-cell imaging

Human embryonic kidney 293T cells were transfected with either N-terminally eGFP-tagged H1.4 (eGFP-H1) alone or in combination with C-terminally mCherry-tagged 70-kDa subunit of the heterotrimeric RPA (RPA-mCherry) and cultured in Dulbecco’s modified Eagle’s medium with 10% fetal bovine serum and 4 mM glutamine at 37 °C in 5% CO_2_. Live-cell imaging was conducted on a Leica SP8 confocal microscope with an inverted stand. For experiments investigating replication stress, cells were subjected to either mock (1% PBS and 0.1% DMSO, v/v) or treatment conditions. Treatment involved exposure to 2 mM hydroxyurea (Sigma-Aldrich) and 20 µM ceralasertib (MedChemExpress) for 12 hours before imaging. At least ten cells were measured for each experimental condition. Puncta were counted using the 3D Object Counter plugin in FIJI (*56*). Images were manually thresholded with a size filter minimum of 10 voxels.

#### Statistical analysis

Errors reported in this study represent the standard error of the mean (SEM). *P* values were determined from two-tailed two-sample *t* tests (ns, not significant; \**P* < 0.05; \*\**P* < 0.01; \*\*\**P* < 0.001).

## Acknowledgments

We thank the O’Donnell laboratory (Rockefeller University) for RPA, Alexey Soshnev (Rockefeller) for helpful discussions, and Murray Tipping and Yevgeniy Romin (MSK Molecular Cytology core facility) for advice with live-cell imaging and analysis. A.O. is supported by the National Science Foundation Graduate Research Fellowship (2016217612). B.Z. is supported by the National Institutes of Health grant R35GM133580. Y.D. is supported by NIH grant R35GM138386, CCSG core grant P30CA008748, the Mr. William H. Goodwin and Mrs. Alice Goodwin and the Commonwealth Foundation for Cancer Research and the Center for Experimental Therapeutics at MSKCC, the Parker Institute for Cancer Immunotherapy and the Anna Fuller Cancer Research Foundation. S.L. is supported by the Robertson Foundation, the Alfred P. Sloan Foundation, and an NIH Director’s New Innovator Award (DP2HG010510). Y.D. and S.L. also acknowledge support from the Pershing Square Sohn Cancer Research Alliance. Y.D. is a Josie Robertson Young Investigator.

## Contributions

R.L. prepared the DNA samples, performed single-molecule binding, *in vitro* phase separation, and droplet fusion experiments with help from G.C.. A.O. prepared the H1 constructs and performed *in vivo* experiments with help from S.C.-R.. J.W. developed analysis software for the single-molecule data. A.L. and B.Z. performed MD simulations. Y.D. and S.L. oversaw the project. All authors contributed to the writing of the manuscript.

