## Supplemental Information for "Single-stranded nucleic acid sensing and coacervation by linker histone H1"

### SUPPLEMENTARY INFORMATION

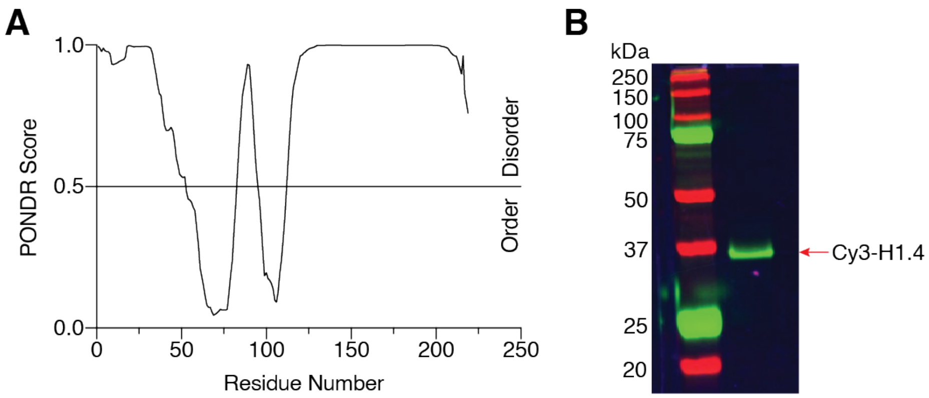


**Supplementary Figure 1. H1 purification and labeling.** (**A**) Predictor of natural disordered regions (PONDR) score for the H1.4 amino acid sequence (generated from www.pondr.com). A score of >0.5 is considered intrinsically disordered. (**B**) SDS-PAGE gel scanned for fluorescence showing purified Cy3-H1.4.

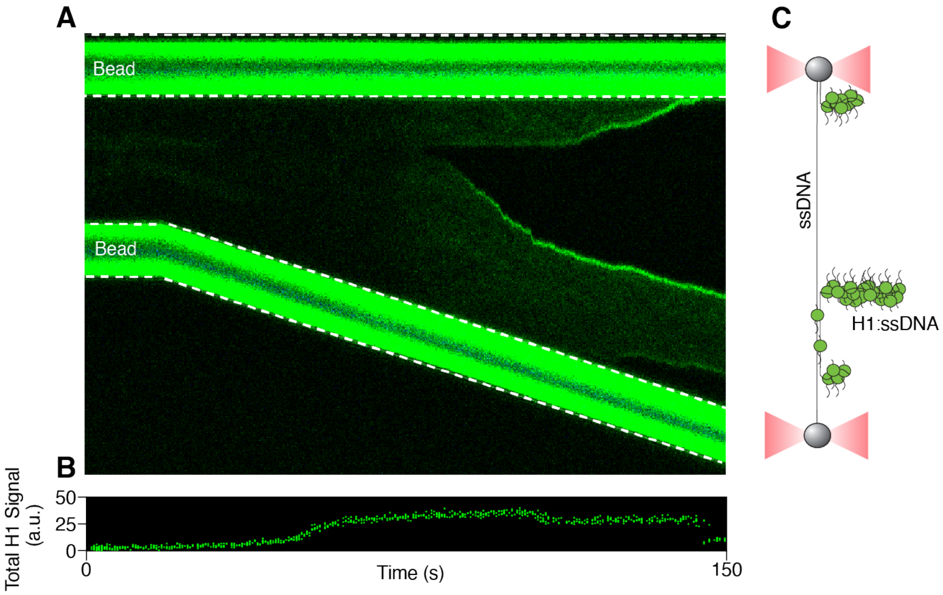


**Supplementary Figure 2. Additional kymograph of H1-DNA interaction.** (**A**) A kymograph showing Cy3-H1 binding to λ-DNA over time as the inter-bead distance was increased. (**B**) Total Cy3 intensity across the DNA tether over time for the kymograph shown in (A). (**C**) Schematic of the final H1 binding configuration for the example shown in (A).

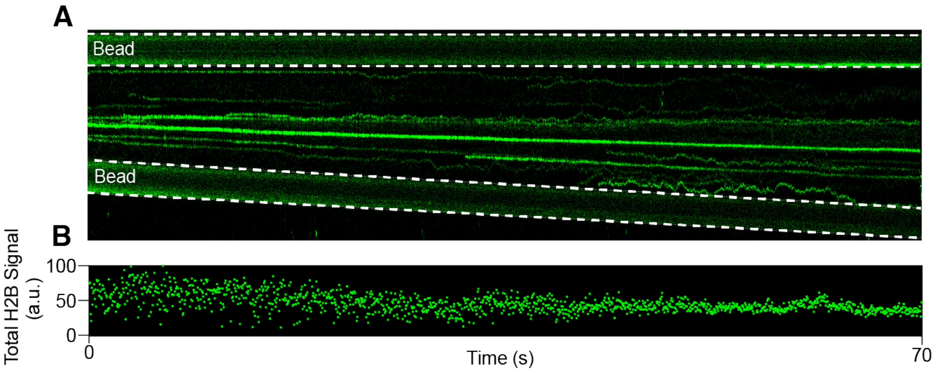


**Supplementary Figure 3. Example kymograph of H2B-DNA interaction.** (**A**) A representative kymograph showing Cy3-H2B binding to λ-DNA over time as the inter-bead distance was increased. (**B**) Total Cy3 intensity across the DNA tether over time for the kymograph shown in (A).

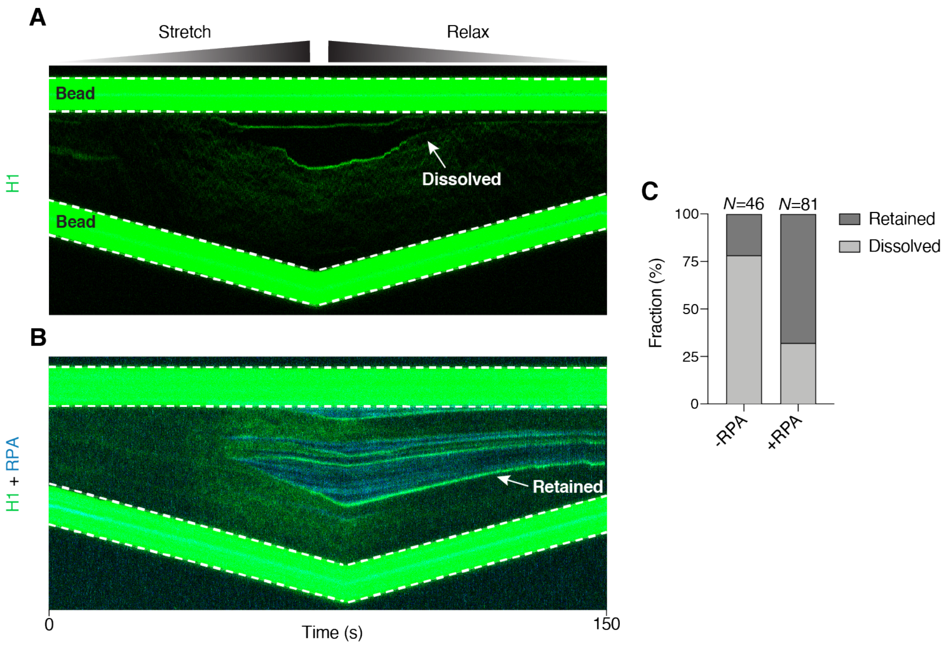


**Supplementary Figure 4. Reversibility of H1:ssDNA foci.** (**A**) A representative kymograph showing reversible formation and dissolution of Cy3-H1 foci during DNA tether stretching and relaxing. (**B**) A representative kymograph showing the persistence of Cy3-H1 foci after tether relaxing in the presence of AlexaFluor488-RPA. (**C**) Fraction of H1 foci dissolved or retained in the absence and presence of RPA.

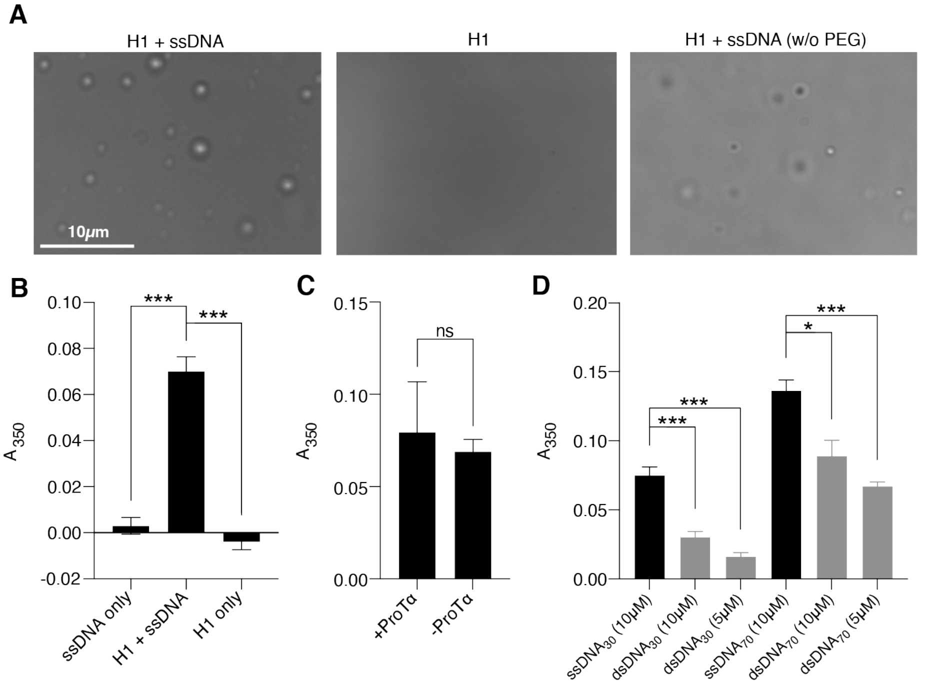


**Supplementary Figure 5. Additional results from bulk phase-seperation assays.** (**A**) Brightfield images of 2.5 μM H1 mixed with 10 μM $ssDNA_{75}$ (left), 2.5 μM H1 alone (middle), and 10 μM H1 mixed with and 10 μM $ssDNA_{75}$ in the absence of PEG (right). (**B**) Solution turbidity ($A_{350}$) measurements for 10 μM $ssDNA_{75}$ alone, 2.5 μM H1 mixed with 10 μM $ssDNA_{75}$, and 2.5 μM H1 alone. (**C**) $A_{350}$ measurements for 2.5 μM H1 mixed with 10 μM $ssDNA_{75}$ in the presence or absence of the H1 chaperone ProTα (2.5 μM). (**D**) $A_{350}$ measurements for 2.5 μM H1 mixed with 10 μM ssDNA, 10 μM dsDNA, or 5 μM dsDNA (to have the same number of nucleotides as 10 μM ssDNA) of 30 or 70 nt/bp in length. Data are mean ± SEM of at least three independent measurements.

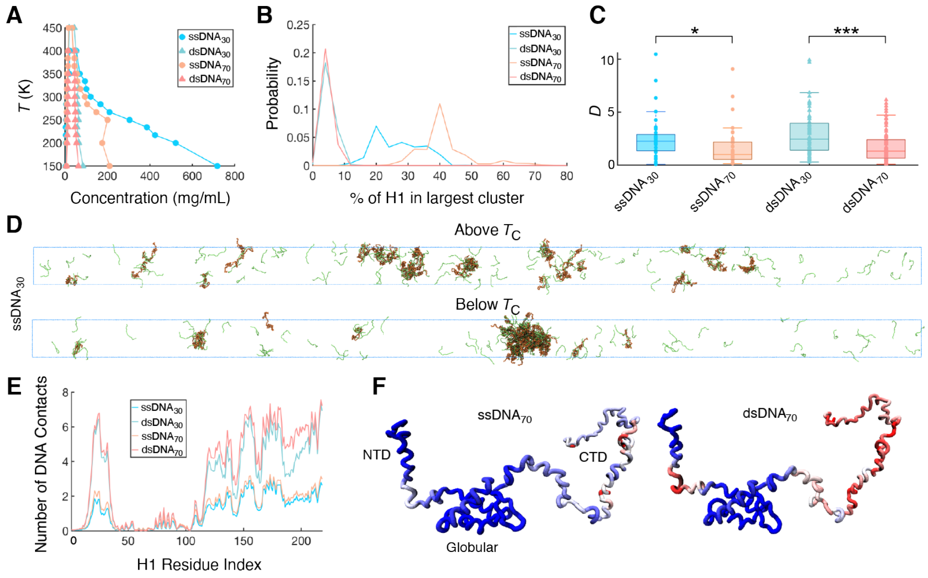


**Supplementary Figure 6. Computational examination of DNA-driven H1 phase separation.** (**A**) Phase behavior of the four simulated systems as a function of temperature. (**B**) Probabililty distribution of the percentage of H1 molecules found in the largest cluster at a temperature of 300 K. (**C**) Diffusion coefficient (*D*) of H1 in each system at 300 K. All values were normalized by the median *D* for $ssDNA_{70}$. (**D**) Representative configurations for the $ssDNA_{30}$ system above and below $T_C$. (**E**) Average number of DNA residues in contact with any given H1 residue, as a function of the H1 residue index. (**F**) Average number of DNA residues in contact with any given H1 residue projected onto a structure of H1. Data ranges from most contacts (red) to least contacts (blue).

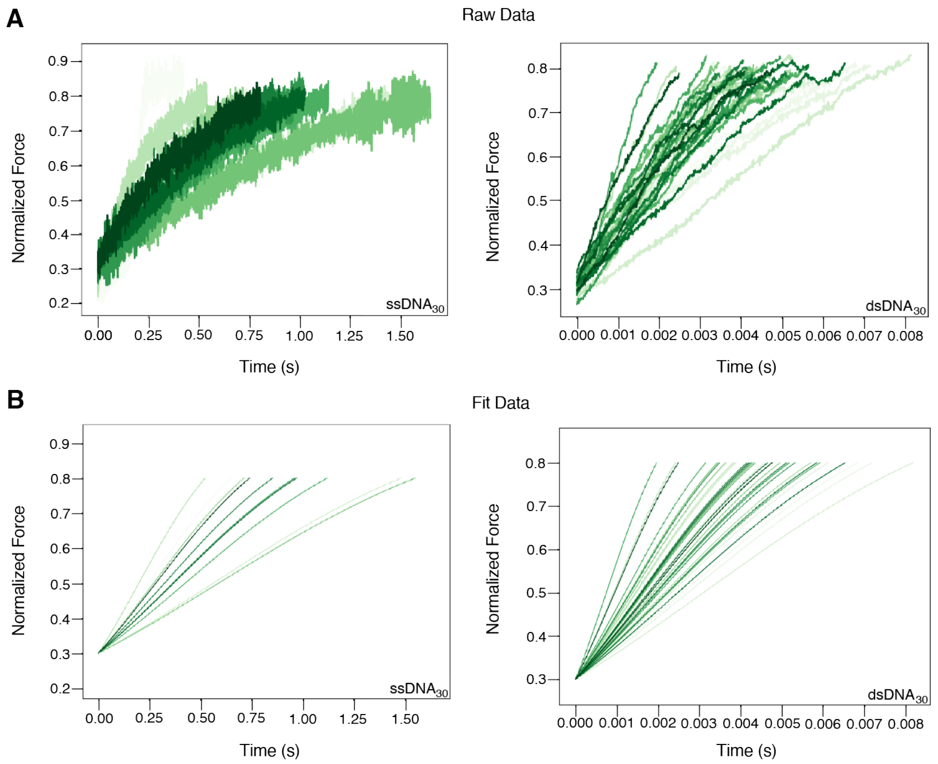


**Supplementary Figure 7. Force profiles from droplet fusion experiments.** (**A**) Raw data between the normalized force values of 0.3 and 0.8 for H1:ssDNA$_{30}$ (left) and H1:dsDNA$_{30}$ (right) droplets. Each trace represents a fusion event between two optically trapped droplets. (**B**) Corresponding sigmoidal fits of the force-time traces shown in (A).

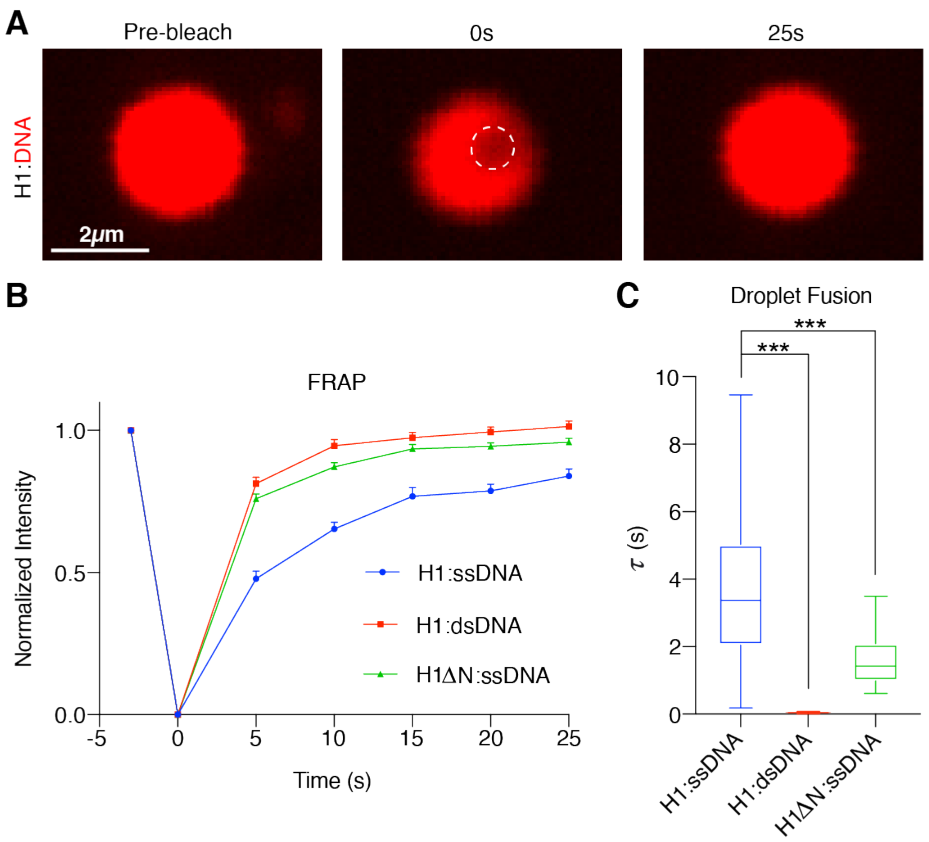


**Supplementary Figure 8. Additional analyses of the biophysical properties of H1:DNA droplets.** (**A**) A representative series of images during the photobleaching and fluorescence recovery of an H1:Cy5-ssDNA$_{75}$ droplet. (**B**) Kinetics of fluorescence recovery for H1:Cy5-ssDNA$_{75}$ (blue), H1:Cy5-dsDNA$_{75}$ (red), and H1ΔN:ssDNA$_{75}$ (green) droplets. At least 15 droplets were used for FRAP analysis for each condition. (**C**) Droplet fusion time ($\tau$) for H1:Cy5-ssDNA$_{75}$, H1:Cy5-dsDNA$_{75}$, and H1ΔN:ssDNA$_{75}$ droplets. At least 12 fusion events were analyzed for each condition. All experiments were performed with 2.5 µM H1 and 10 µM DNA (10% labeled). Data are mean ± SEM.

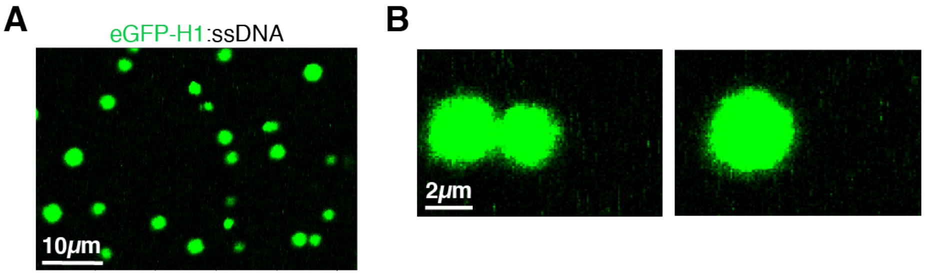


**Supplementary Figure 9. eGFP-H1:ssDNA droplet formation and fusion *in vitro*.** (**A**) A representative image of droplets formed with 2.5 μM recombinant eGFP-H1 and 10 μM $ssDNA_{75}$ visualized by GFP fluorescence. (**B**) Snapshots of two eGFP-H1:$ssDNA_{75}$ droplets pre- and post-fusion.

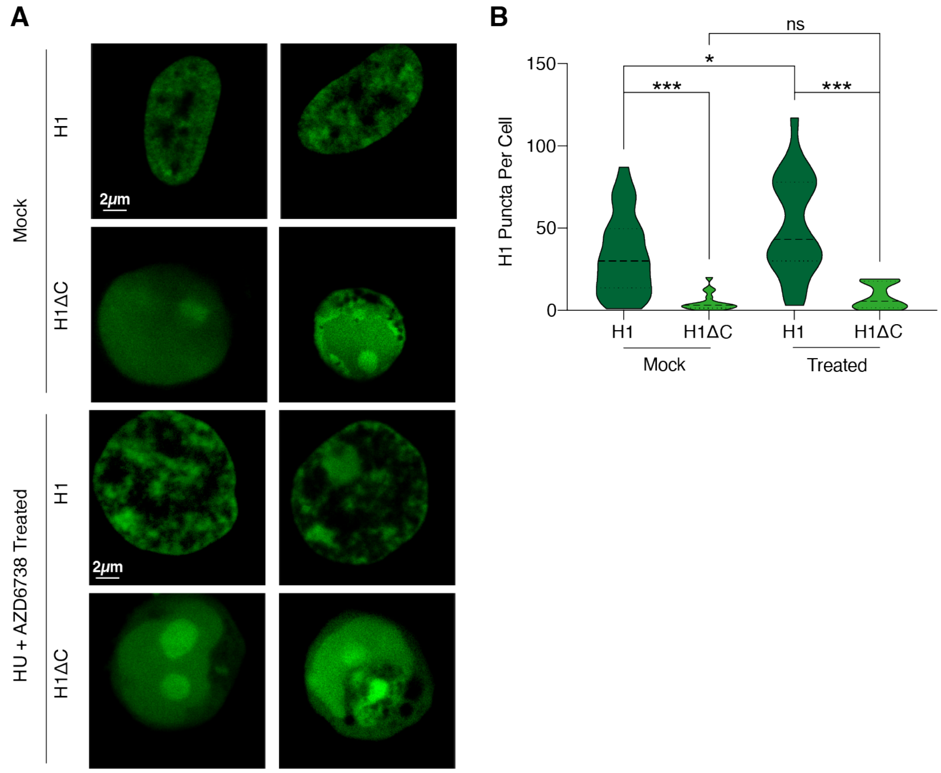


**Supplementary Figure 10. Additional live-cell imaging results.** (**A**) Representative confocal fluorescence images of HEK293T cells transfected with either eGFP-H1 or eGFP-H1ΔC and treated with either mock or 2 mM HU + 20 μM AZD6738 for 12 h. (**B**) Violin plot showing the distribution of number of H1 puncta per cell for eGFP-H1 and eGFP-H1ΔC cells after mock or HU + AZD6738 treatment.
